# Artifact-free whole-slide imaging with structured illumination microscopy and Bayesian image reconstruction

**DOI:** 10.1101/765396

**Authors:** Karl Johnson, Guy M. Hagen

## Abstract

**Background:** Structured illumination microscopy (SIM) is a method which can be used to image biological samples and can achieve both optical sectioning and super-resolution effects. Optimization of the imaging setup and data processing methods results in high quality images without artifacts due to mosaicking or due to the use of SIM methods. Reconstruction methods based on Bayesian estimation can be used to produce images with a resolution beyond that dictated by the optical system.

**Findings:** Five complete datasets are presented including large panoramic SIM images of human tissues in pathophysiological conditions. Cancers of the prostate, skin, ovary, and breast, as well as tuberculosis of the lung, were imaged using SIM. The samples are available commercially and are standard histological preparations stained with hematoxylin and eosin.

**Conclusion:** The use of fluorescence microscopy is increasing in histopathology. There is a need for methods which reduce artifacts when employing image stitching methods or optical sectioning methods such as SIM. Stitched SIM images produce results which may be useful for intraoperative histology. Releasing high quality, full slide images and related data will aid researchers in furthering the field of fluorescent histopathology.

## Data description

### Context

Structured illumination microscopy (SIM) is a method in optical fluorescence microscopy which can achieve both optical sectioning (OS-SIM) [1] and resolution beyond the diffraction limit (SR-SIM) [2,3]. SIM has been used for super-resolution imaging of both fixed and live cells [4–7] and has matured enough as a method that it is now available commercially. In SIM, a set of images is acquired using an illumination pattern which shifts between each image. As SIM has developed, diverse strategies have been proposed for creation of the SIM pattern [1,8–13]. Several different approaches for processing the data have also been introduced [3,7,8,14–16].

Recently, microscope systems capable of imaging with high resolution and a large field of view (FOV) have been developed [17–21], some using custom-made microscope objectives. However, stitching together images acquired with a higher magnification objective to create a large mosaic remains a valid and popular approach. Some published results involving stitched images suffer from pronounced artifacts in which the edges of the individual sub-images are visible, usually as dark bands which outline each sub-image [22–24]. On the other hand, several studies have proposed methods for stitching of microscope images with reduced artifacts [25–32].

The combination of SIM with image stitching methods allows collection of large FOV images with both optical sectioning and super-resolution properties. Here, we demonstrate methods and provide complete datasets for five different samples. The samples are hematoxylin and eosin (H&E) stained histological specimens which provide examples of human diseases (ovarian cancer, breast cancer, prostate cancer, skin cancer, and tuberculosis), and which are also available commercially for those who wish to reproduce our work. We used freely available optical designs [6,10,33] and open source software [33] for SIM imaging, along with freely available software for image stitching (Microsoft Image Composite Editor (ICE) [34], or a well validated plugin [26] for ImageJ [35]). Combining this with devignetting methods, we produced stitched images which are free of noticeable artifacts from stitching or from SIM reconstruction.

Fluorescence microscopy is becoming more important in histopathology. Traditional bright field microscopy diagnostic methods require a time-consuming process, involving chemical fixation and physical sectioning. The use of optical sectioning fluorescence microscopy allows high-quality images to be captured without the need for physical sectioning. Consequently, it has been shown that imaging can be performed on large human tissue samples within 1 hour after excision [36]. Additionally, other studies have shown the results of fluorescence imaging to be usable and accurate in diagnosis of various medical conditions [37–42]. Previously, it was noted that obvious stitching artifacts significantly decrease the usability of large fluorescence images in medical diagnosis. In one case, such artifacts resulted in the rejection of over half of the images acquired [38]. The setup we describe here allows for fast, artifact-free, high-resolution imaging of fluorescent samples, and is compatible with samples stained with most fluorescent dyes.

## Methods

### Samples

All samples used in this study are available from Carolina Biological, Omano, or Ward’s Science. The samples are approximately 7 µm thick and are stained with hematoxylin and eosin. The commercial source, product number, and other SIM imaging parameters for each sample are detailed in Table 1. Table 2 details imaging parameters for acquisitions of each sample with a color camera.

**Table 1:** Imaging parameters for the SIM datasets.

| Sample | Source company and product no. | SIM pattern no. of phases | Exposure time, ms | No. of tiles | Objective mag/NA | Acquisition time, s | Stitching software |
| --- | --- | --- | --- | --- | --- | --- | --- |
| Carcinoma of Prostate | Carolina, 318492 | 5 | 50 | 23 × 11 | 20×/0.45 | 315 | Microsoft ICE |
| Basal Cell Carcinoma | Ward’s Science, 470183-256 | 6 | 75 | 29 × 18 | 30×/1.05 | 821 | FIJI |
| Adenocarcinoma of Ovary | Carolina, 318628 | 5 | 100 | 25 × 14 | 10×/0.4 | 595 | Microsoft ICE |
| Adenocarcinoma of Breast | Carolina, 318766 | 8 | 200 | 12 × 8 | 10×/0.4 | 278 | FIJI |
| Lung Tuberculosis | Omano, OMSK-HP50 | 5 | 100 | 20 × 16 | 30×/1.05 | 541 | FIJI |

**Table 2:** Parameters for the color images.

| Sample | No. of tiles | Objective mag/NA |
| --- | --- | --- |
| Carcinoma of Prostate | 6 × 5 | 4×/0.16 |
| Basal Cell Carcinoma | 5× 5 | 4×/0.16 |
| Adenocarcinoma of Ovary | 11 × 11 | 4×/0.16 |
| Adenocarcinoma of Breast | 6 × 6 | 4×/0.16 |
| Lung Tuberculosis | 8 × 10 | 10×/0.4 |

### Microscope setup and data acquisition

We used a home-built SIM setup based on the same design as described previously [6,10,15] (Fig. 1). The SIM system is based on an IX83 microscope (Olympus) equipped with a Zyla 4.2+ sCMOS camera (Andor) under the control of IQ3 software (Andor). We used the following Olympus objectives: UPLSAPO 4×/0.16 NA, UPLSAPO 10×/0.4 NA, LUCPLFLN 20×/0.45 NA, and UPLSAPO 30×/1.05 NA silicone oil immersion. For color images we used an aca1920-40uc color camera (Basler) under control of Pylon software (Basler). We used a MS-2000 motorized microscope stage (Applied Scientific Instrumentation) to acquire tiled SIM images. In all datasets, the stage scanning was configured such that all image edges overlapped by 20%.

**Figure 1:**
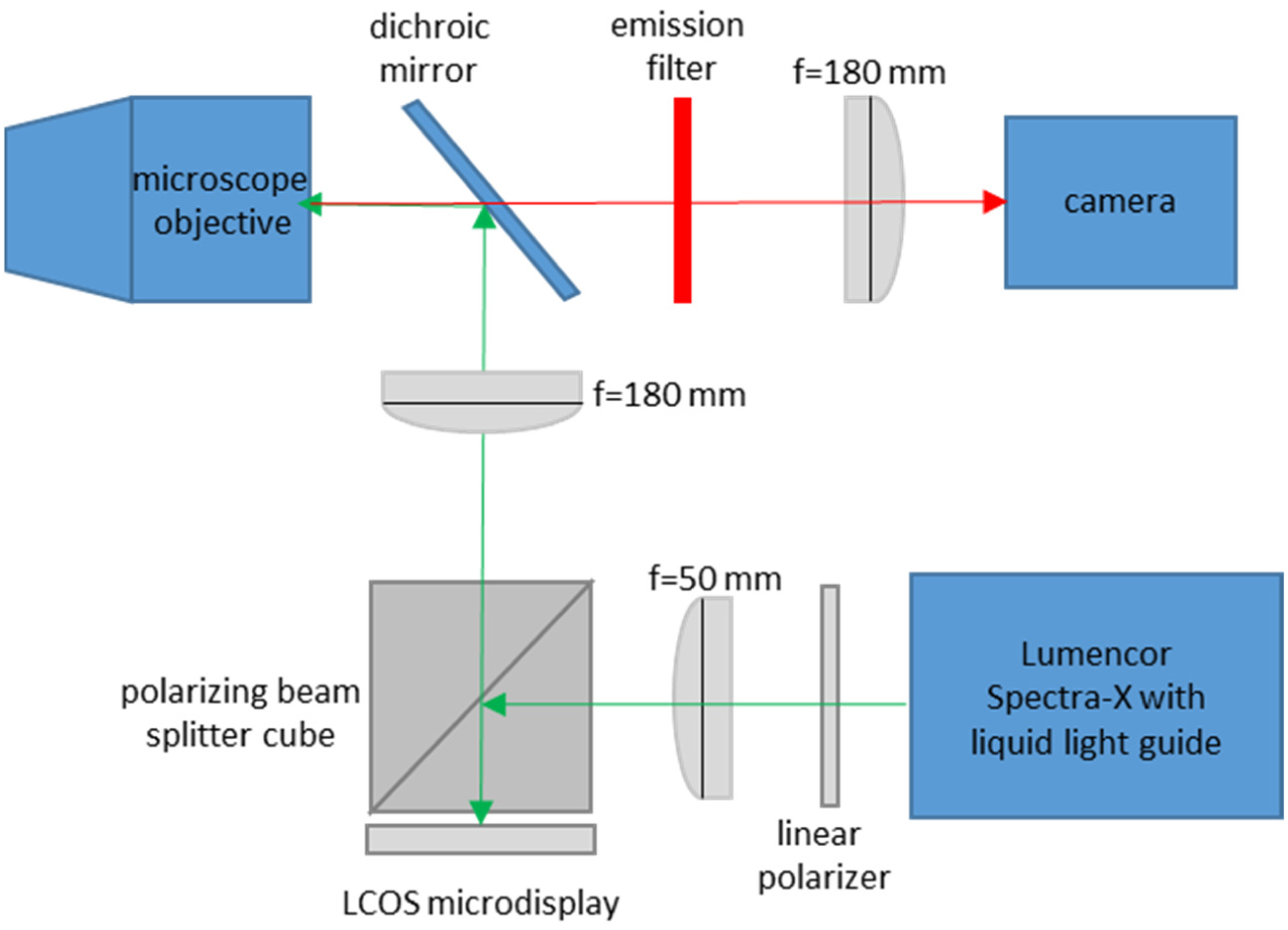
Simplified diagram of SIM system. LCOS, liquid crystal on silicon.

Briefly, the SIM system uses a ferroelectric liquid crystal on silicon (LCOS) microdisplay (type SXGA-3DM, Forth Dimension Displays). This device has been used previously in SIM and related methods in fluorescence microscopy [5,10,15,33,43–47] and allows one to produce patterns of illumination on the sample which can be reconfigured at will by changing the image displayed on the device. The light source (Lumencor Spectra-X) is toggled off between SIM patterns and during camera readout. Close synchronization between the camera acquisitions, light source, and microdisplay ensures rapid image acquisition, helps reduce artifacts, and reduces light exposure to the sample. The supplementary material contains more information about system integration.

### SIM data processing

SIM reconstructions were performed using SIMToolbox, an open-source and freely available program that our group developed for processing SIM data [33]. We generated optically sectioned, enhanced resolution images using a Bayesian estimation method, maximum *a posteriori* probability SIM (MAP-SIM) [15]. MAP-SIM works using maximum *a posteriori* probability methods, which are well known in microscopy applications [48,49], to enhance high spatial frequency image information. We then combine this information, in the frequency domain, with low spatial frequency image information obtained by OS-SIM methods, then produce the final image by an inverse Fourier transform [15]. We typically measure the final resolution obtained by analyzing the frequency spectrum of the resulting image, as is discussed below.

The illumination patterns used here are generated such that the sum of all positions in each pattern set results in homogenous illumination. As such, a widefield (WF) image can be reconstructed from SIM data simply by performing an average intensity projection of the patterned images. This can be described by

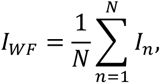

where N is the number of pattern phases, I_n_ is the image acquired on the n^th^ illumination position, and I_WF_ is the WF reconstruction. This is the method we used to generate WF images throughout this study.

### Vignetting correction

Following SIM reconstruction, vignetting artifacts remain in each tile. If not removed prior to stitching, this vignetting introduces a distracting grid pattern in the final stitched image. We performed vignette removal by dividing each tile of the mosaic by an image representing the vignetting profile common to all tiles. Other studies have used an image of a uniformly fluorescent calibration slide as a reference for vignette removal [36], where information concerning non-uniform illumination is captured. However, we found that SIM processing introduces vignetting artifacts beyond those due to non-uniform illumination. Additionally, these artifacts vary depending on properties of the sample being imaged. As such, performing pre-acquisition calibration on a uniformly fluorescent slide is not sufficient to remove vignetting artifacts from SIM reconstructions. Instead, an estimate of the vignetting profile is found through analysis of the mosaic tiles after SIM reconstruction.

A blurred average intensity projection of the tiles is a good approximation of the vignetting profile, as an average intensity projection merges the tiles into a single image with reduced foreground information while preserving vignetting. Subsequent blurring with an appropriate radius and edge-handling method also eliminates the high spatial frequency foreground without impacting the low spatial frequency illumination profile. To eliminate errors during the blurring step due to the blurring area extending outside the original image, we used an edge handling method in which the blurring area is reduced near the edges of the image such that no values outside the image border are sampled. Unlike edge handling methods in which the image is padded with a uniform value (or mirrored and tiled) to accommodate a blurring area which extends beyond the original image limits, this method is free from major artifacts, such as erroneous brightness of the image edges (see supplementary figure S1). This approximation of the illumination profile works especially well for histological samples, as such samples are non-sparse and require many tiles, factors which improve the accuracy this approach. We performed all steps of this devignetting process using built-in functions and the ‘Fast Filters’ plugin in ImageJ [50]. The effect of devignetting is illustrated in Fig. 2.

**Figure 2:**
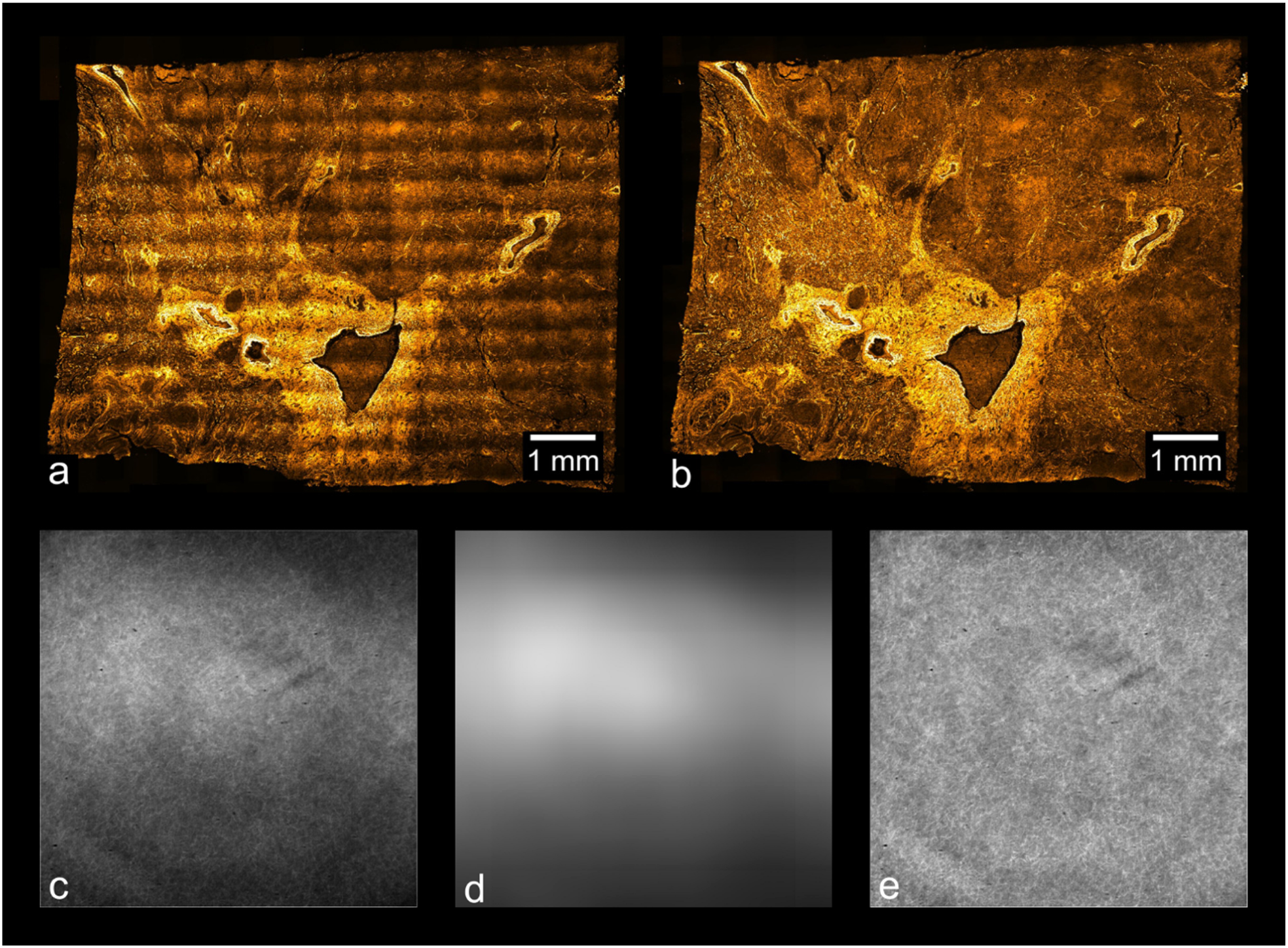
Vignetting artifacts and their removal. (a) shows the result of stitching images without applying the devignetting process, while (b) shows a stitch of the same data after devignetting has been applied. (c) shows the average intensity projection of the images used to stitch (a), which estimates the vignette profile of each frame. This estimate can be refined by application of an edge-limited blurring filter, as shown in (d). (e) shows the average intensity projection of the data used in (b), after devignetting has been applied. The uniform brightness of (e) indicates that no major vignetting artifacts remain in the devignetted data.

### Image Stitching

With visible vignetting removed, we then stitched together a composite image from the tiles. The pre-processing allows for stitching to be done in various stitching applications; Microsoft ICE and Preibisch’s plugin for FIJI [26] were used to stitch the data presented here.

The data processing procedure is summarized in Fig. 3. The total time for processing each dataset was about 30 min.

**Figure 3.**
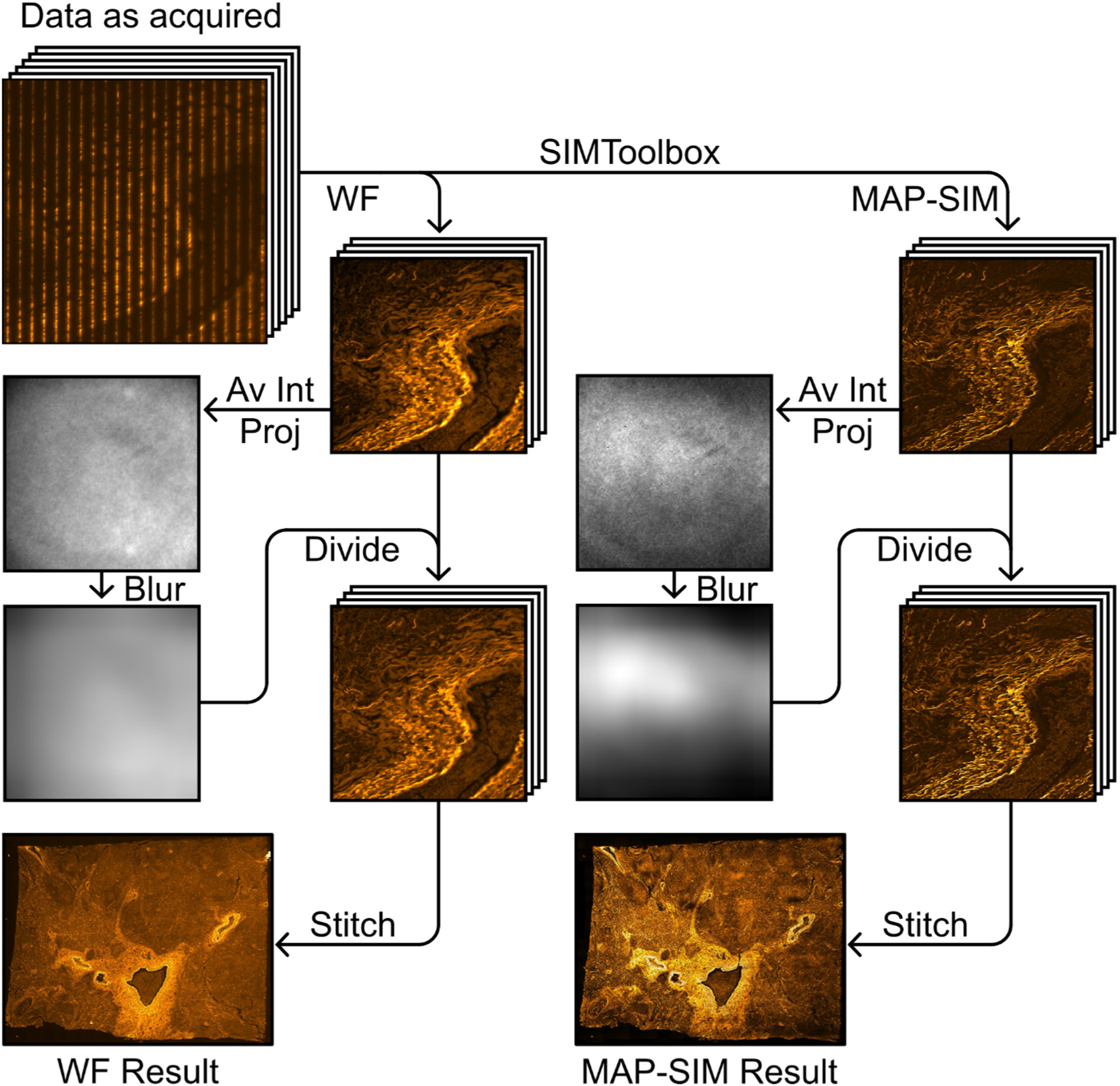
Panoramic SIM data processing workflow. Devignetting was performed after SIM reconstruction. Note the vignette profile differs between reconstruction methods, necessitating separate projection, blurring and division steps. Av Int Proj refers to average intensity projection.

### Color image data processing methods

We created color overview images by stitching devignetted brightfield acquisitions. Devignetting was performed simply by adding the inverse of an empty brightfield acquisition to each color tile using ImageJ. For this method to produce optimal results, the empty brightfield image must be acquired in conditions identical to those of the raw tile data, such that the illumination profile in the empty image matches that of the unprocessed tiles. This simple operation removes nearly all visible vignetting and color balance artifacts within each tile. The results after devignetting were then stitched using Preibisch’s plugin for FIJI [26].

### Resolution measurement

We evaluated our results by measuring image resolution using SR Measure Toolbox. SR Measure Toolbox [51] measures the resolution limit of input images through analysis of the normalized, radially averaged power spectral density (PSD_ca_) of the images, as previously described [6]. Briefly, the resolution limit in real space is determined by evaluating the cutoff frequency in Fourier space. The cutoff frequency is estimated by calculating the spatial frequency at which the PSD_ca_ (after noise correction) drops to zero.

Focusing on the basal cell carcinoma sample, we selected 125 (out of 522 total) image tiles, calculated the PSD and resolution for each tile, and averaged the results. We found that, in the case of this sample, the image resolution was 593 ± 20 nm for WF and 468 ± 2.5 nm for MAP-SIM (average ± standard deviation). This data was acquired with a UPLSAPO 30×/1.05 NA silicone oil immersion objective. Figure 4 shows an example measurement for one image tile. Figure 5 shows a plot of PSD_ca_ for this image tile.

**Figure 4.**
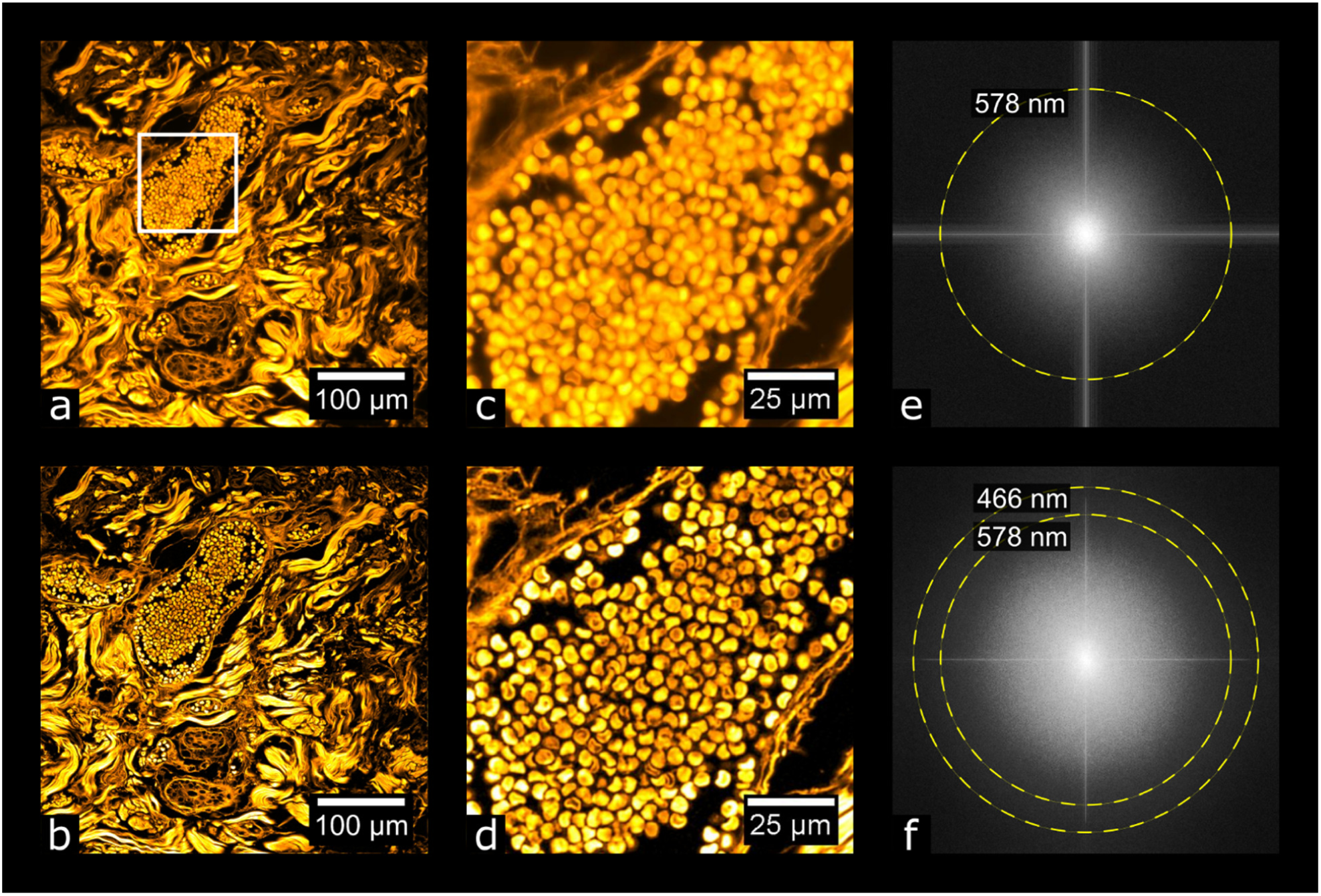
Evaluating image resolution. (a) and (b) show a tile from the data in Fig. 7 (basal cell carcinoma sample) after widefield and MAP-SIM reconstruction, respectively. (c) and (d) each show a zoom-in of (a) and (b), respectively. (e) and (f) each show the FFT of (a) and (b), respectively. The dotted lines in (e) and (f) indicate the resolution of each image according to the resolution measurement described.

**Figure 5.**
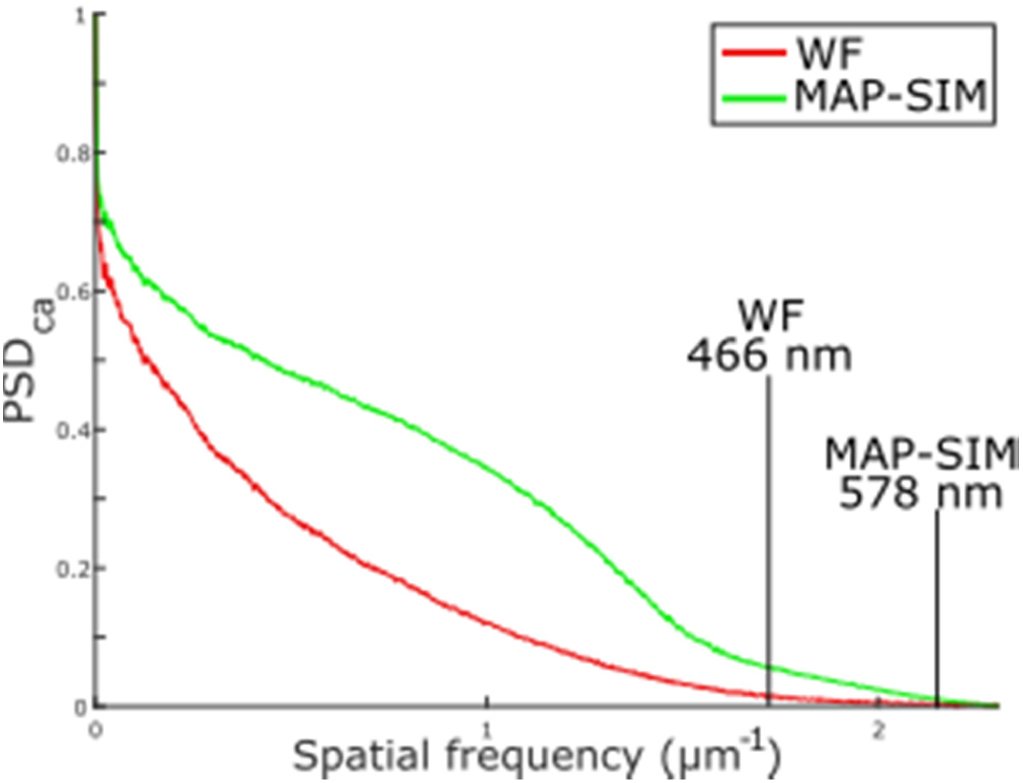
Normalized, radially averaged power spectral density (PSD_ca_) and resolution analysis measured on the tiles shown in Figs. 4a and 4b.

## Results

Figure 6 shows images of a prepared slide containing a human prostate carcinoma sample stained with H&E. Fig. 6a shows a stitched color overview, and Fig. 6d shows a zoom-in of the region indicated in Fig. 6a, acquired separately using a UPLSAPO 20×/0.75NA objective. Fig. 6b shows a stitched widefield fluorescence image, and Fig. 6c shows a stitched SIM image. Figs. 6e and 6f each show zoom-ins of the stitches shown in Figs. 6b and 6c, respectively. Using the acquisition and processing methods described, whole-slide images are produced without any visible stitching artifacts. Additionally, the MAP-SIM reconstruction method produces resolution superior to that of the widefield data.

**Figure 6:**
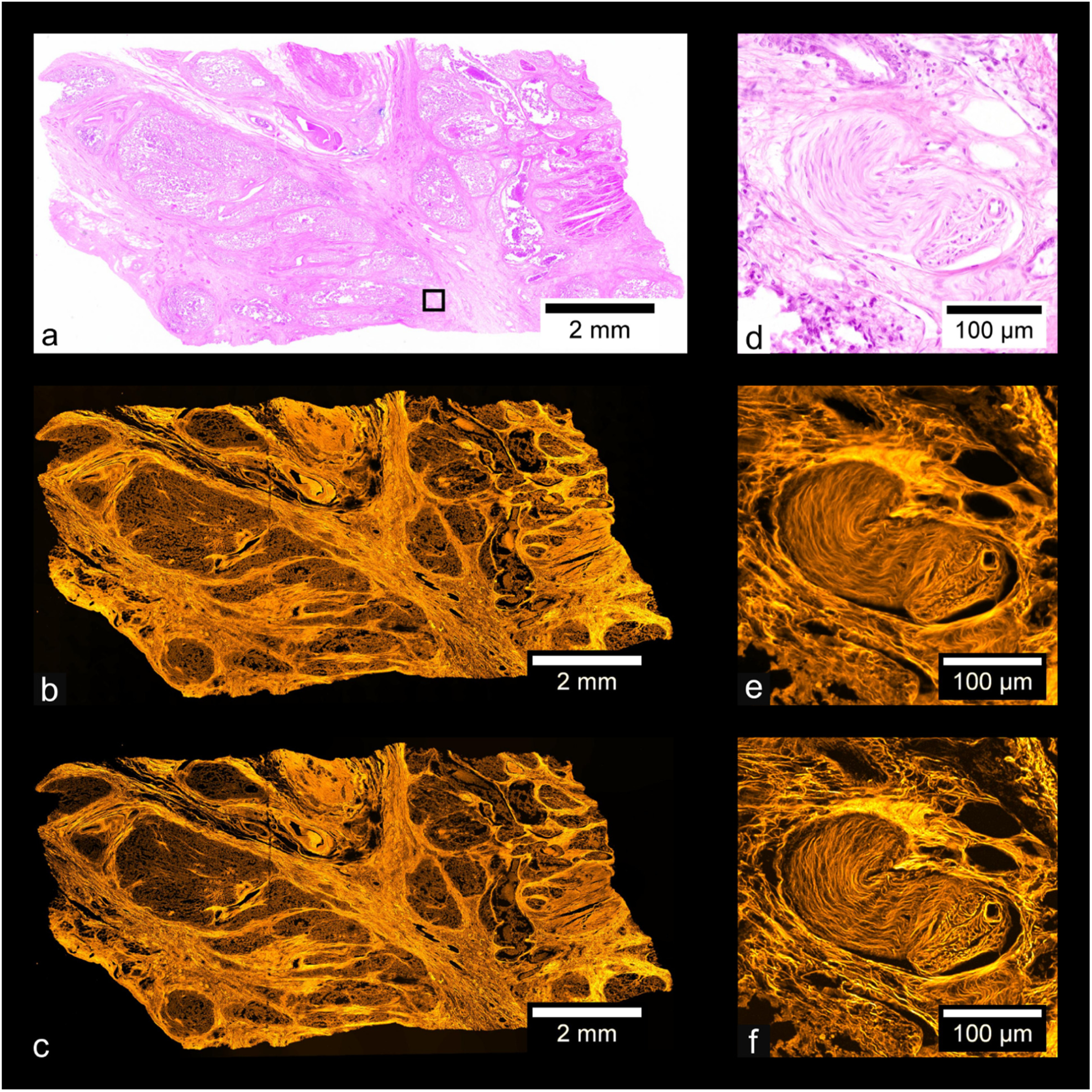
Carcinoma of human prostate. (a) Color overview, (b) WF stitch, (c) MAP-SIM stich. (d) shows a region of the sample indicated in (a). (e) and (f) each show a zoom-in of (b) and (c), respectively, in the region indicated in (a).

Figures 7-10 show similar comparisons for basal cell carcinoma, ovary adenocarcinoma, breast adenocarcinoma, and tuberculosis of the lung, respectively.

**Figure 7:**
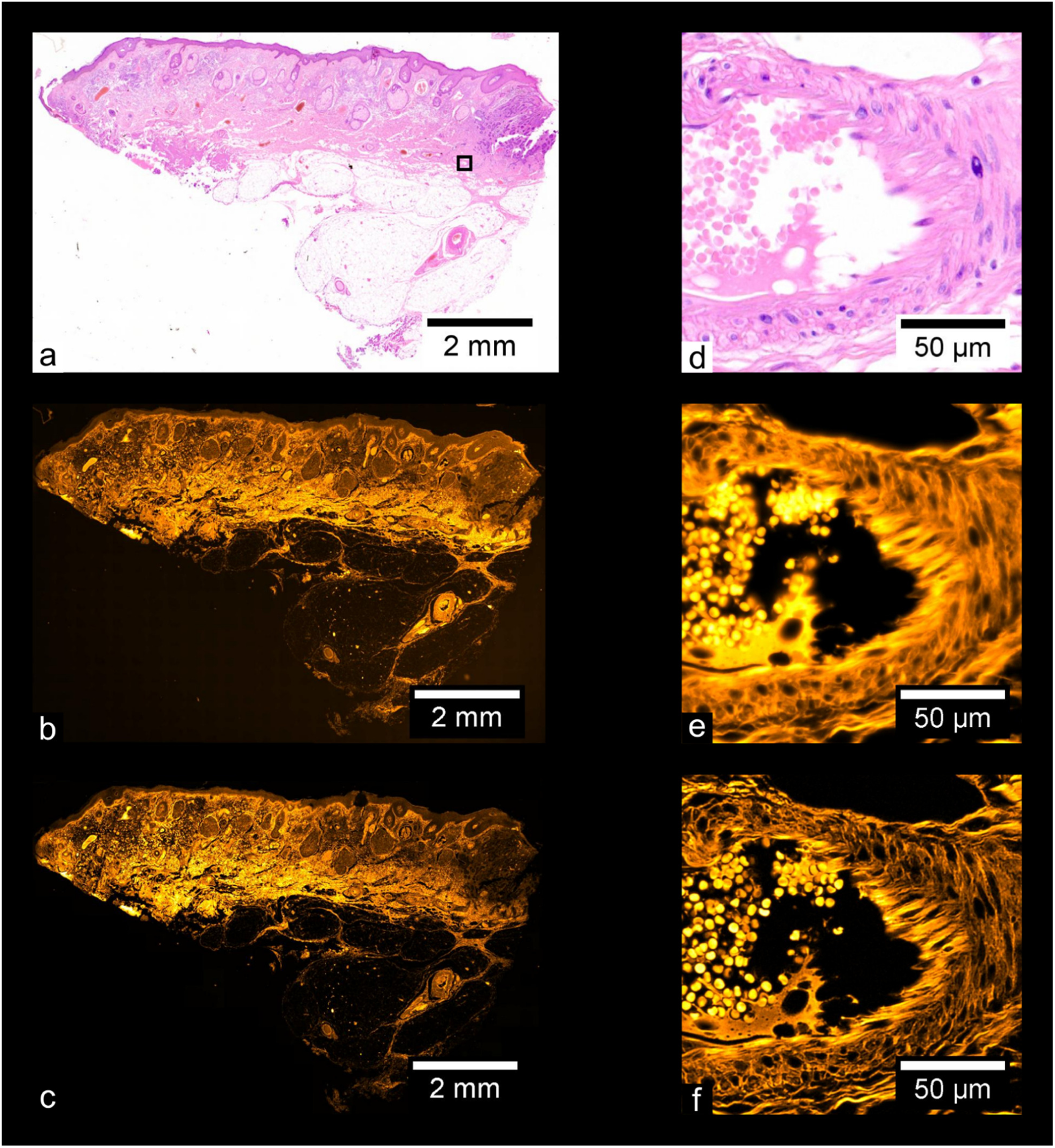
Basal Cell Carcinoma. (a) Color overview, (b) WF stitch, (c) MAP-SIM stich. (d) shows a region of the sample indicated in (a). (e) and (f) each show a zoom-in of (b) and (c), respectively, in the region indicated in (a).

**Figure 8:**
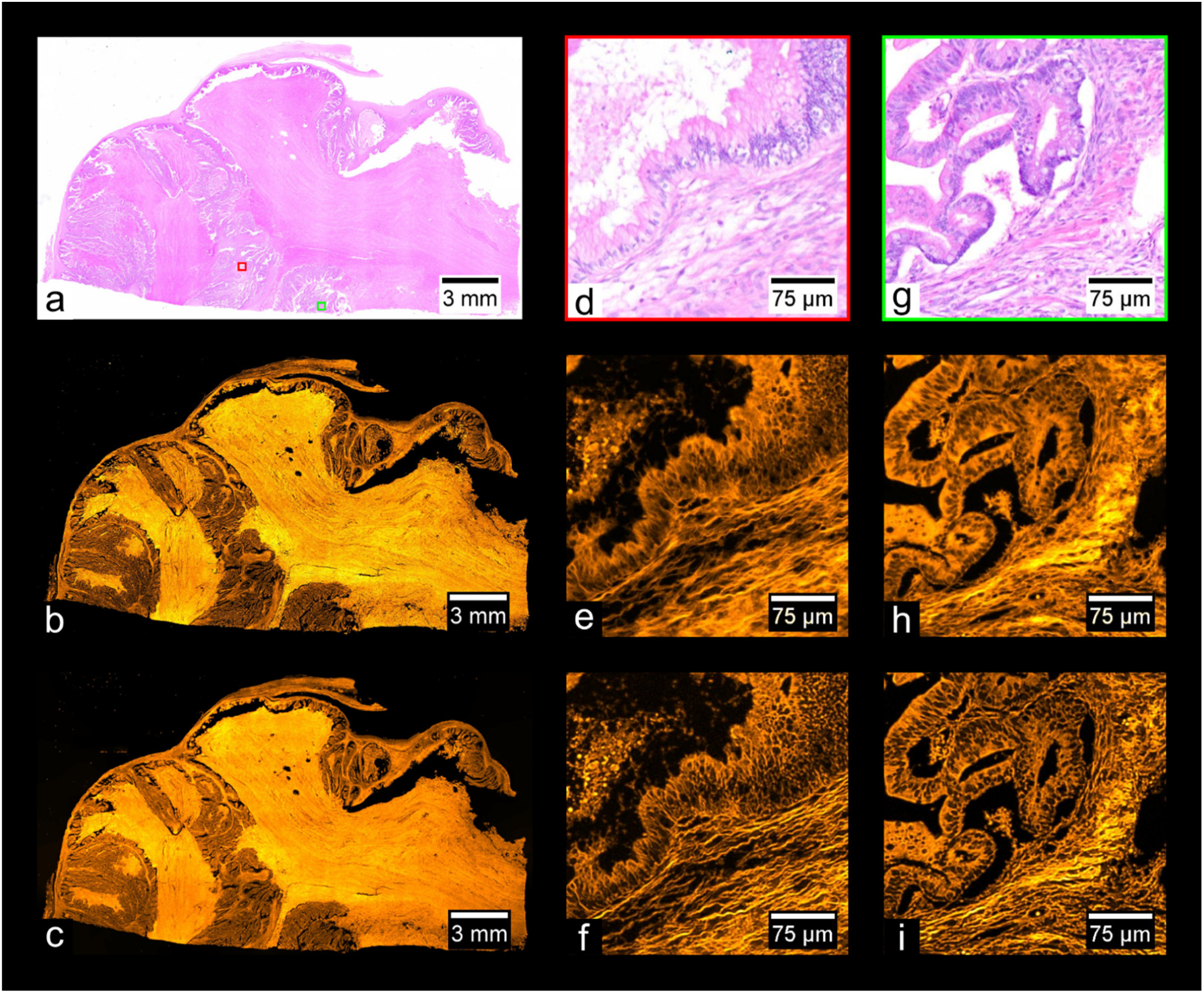
Adenocarcinoma of human ovary. (a) Color overview, (b) WF stitch, (c) MAP-SIM stich. (d), (g) show a region of the sample indicated in (a), acquired separately from (a) using a 10× objective. (e) and (h) show a zoom-in of (b), while (f) and (i) show a zoom-in of (c), all in the regions indicated in (a).

**Figure 9:**
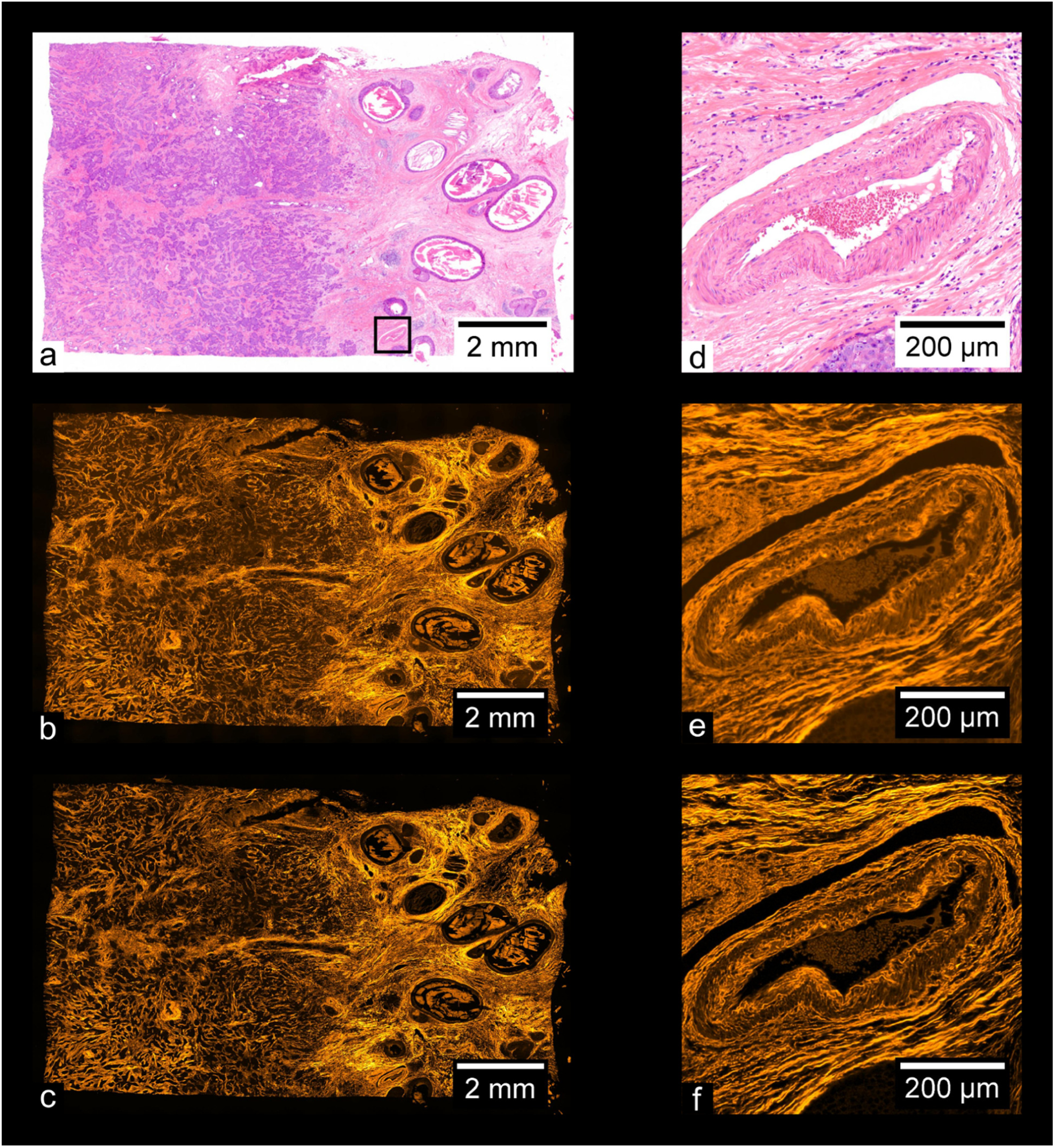
Adenocarcinoma of human breast. (a) Color overview, (b) WF stitch, (c) MAP-SIM stich. (d) shows a region of the sample indicated in (a), acquired separately from (a) using a 10× objective. (e) and (f) each show a zoom-in of (b) and (c), respectively, in the region indicated in (a).

**Figure 10:**
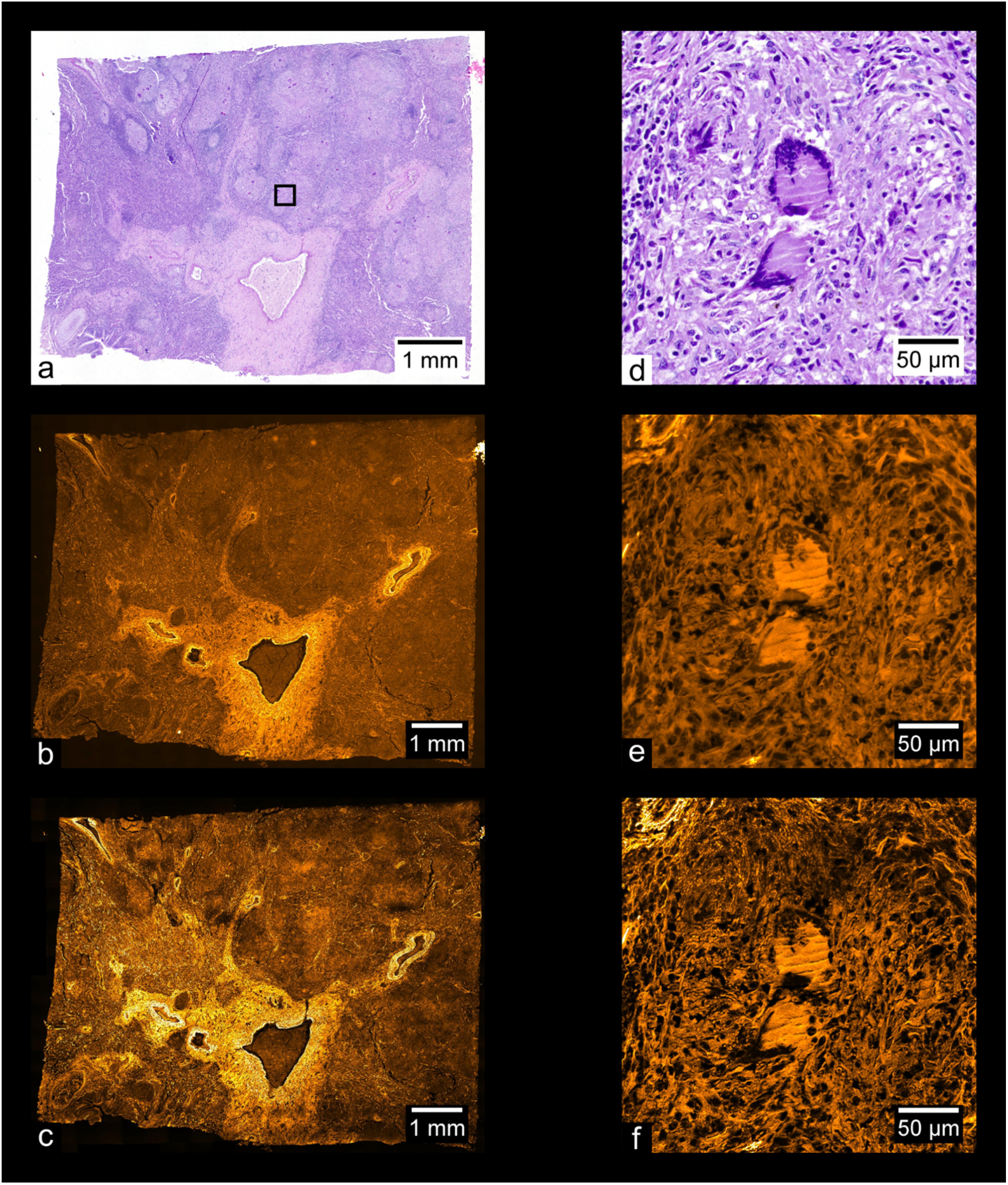
Tuberculosis of human lung. (a) Color overview, (b) WF stitch, (c) MAP-SIM stich. (d) shows a region of the sample indicated in (a), acquired separately from (a) using a 20× objective. (e) and (f) each show a zoom-in of (b) and (c), respectively, in the region indicated in (a).

The data shown in figures 6-10 is freely available through Giga DB [reference to be added]. This dataset includes all color overviews as well as WF and MAP-SIM stitches at full resolution. In addition, all image tiles (prior to devignetting) used to create the WF and MAP-SIM stitches of the basal cell carcinoma sample are provided.

## Discussion

Many past studies into stitching of SIM mosaics have suffered from noticeable image artifacts, arising from flaws in the optical setups used as well as imperfections in the SIM reconstruction and image stitching processes. While these artifacts are sometimes minimal enough to remain uncorrected, certain artifacts seriously inhibit the usefulness of the final stitched image. In [23], the authors note that issues in triggering and evenly illuminating the microdisplay being used for illumination resulted in striping and vignetting artifacts; similarly, in [22,24,36,52], stitching artifacts are apparent in the images. Here, optimization of the optical setup, camera-microdisplay synchronization, and image processing methods yielded whole-slide images free from visible SIM or image stitching artifacts. In addition to the elimination of artifacts, our use of SIMToolbox to perform SIM reconstruction on the data allows for a variety of reconstruction algorithms to be used, including super-resolution algorithms such as MAP-SIM. This too presents an improvement over previous works. Our methods also allow for stitching of high-magnification tiles into large-FOV images with subdiffractive detail (see supplementary Fig. S3).

Another advantage of the acquisition and processing methods demonstrated here is the minimization of user intervention, and in turn, reductions in acquisition and processing time. Firstly, the use of Andor IQ during acquisition allows for stage movement, sample focusing, image acquisition, and SIM pattern advancement to be controlled automatically. Loading of the sample, definition of the mosaic edges, and manual focus on 3-5 positions of the sample are the only steps needed to be taken by the user before acquisition can begin. Recent developments in autofocus technology for SIM may allow for the manual focus step to be shortened or omitted [52]. These automated steps during acquisition allow for large mosaics to be acquired. The quality of the final stitched images does not degrade for larger mosaics – in fact, the quality of the devignetting process improves with larger datasets, as more data is available to produce an accurate estimation of the illumination profile. SIMToolbox (version 2.0), which is capable of utilizing the processing power of modern consumer graphics cards during MAP-SIM processing, also reduces the time spent during the data processing phase. Finally, unlike other super-resolution reconstruction methods such as SR-SIM, MAP-SIM is able to produce artifact-free results without tuning of reconstruction parameters by the user, a process which is difficult to automate and requires significant user experience.

One drawback the method presented here is the inability to image the entire volume of samples thicker than ∼0.5 mm. However, this limitation does not prevent large, unsectioned samples from being imaged, as is the case with bright field microscopy, where samples must be thin enough for transmitted light to reach the objective. Rather, as the light which illuminates the sample in fluorescence microscopy emanates from the objective, all surface regions of a large sample may be imaged. Additionally, due to the optical sectioning exhibited by SIM, light from out-of-focus regions of the sample is almost completely attenuated. Consequently, imaging the surfaces of large samples with SIM produces high-contrast images of thin regions without the need for physical sectioning, as previously demonstrated [23,36].

Here, we demonstrated our imaging techniques on traditionally prepared histopathological samples in order to provide a comparison between bright field imaging and SIM, but the same techniques can be used to image a wide variety of fluorescently labelled samples, as demonstrated in the supplementary material. The ability to seamlessly image the entire surface region of large samples has multiple potential applications in histopathology. SIM presents unique advantages in analyzing the surgical margins of large tissue excisions, as demonstrated by Wang [36]. Briefly, due to the ability of SIM to image an unsectioned sample, analysis of surgical margins using SIM requires imaging of far less surface area than that needed for bright field imaging. Confocal imaging of core needle biopsy samples has been previously demonstrated to produce images suitable for medical diagnosis [42], a practice easily adapted to SIM. The speed at which sample preparation and image acquisition can be performed in fluorescence microscopy presents opportunities for intra-operative analysis of tissue samples using SIM techniques, as mentioned by multiple other studies [23,36,53,54].

### Reuse potential

The data provided here presents various opportunities for reuse. Firstly, the unstitched image tiles provided in the dataset, which still contain vignetting artifacts, may be used to reproduce the results of our devignetting process, as well as to further develop more sophisticated devignetting approaches suited for SIM. These tiles might also be used to create or modify existing stitching software for global minimization of stitching artifacts. For example, the frequency-domain detection of periodic stitching artifacts discussed in the supplementary material could be used to minimize such artifacts in developing new stitching software. With the multiple high-resolution color overviews and stitched SIM images, comparison of structures visible in the brightfield and fluorescent images could be performed to further study the use of fluorescence microscopy in histopathology.

## Availability of source code and requirements

Project name: SIMToolbox version 2.12

Project home page: http://mmtg.fel.cvut.cz/SIMToolbox/

Operating system: platform independent

Programming language: MATLAB

License: GNU General Public License v3.0

## Detailed software compatibility notes

The SIMToolbox GUI was compiled with MATLAB 2015a and tested in Windows 7 and 8. The GUI is a stand-alone program and does not require MATLAB to be installed. To use the MATLAB functions within SIMToolbox (i.e., without the GUI), MATLAB must be installed. The functions were mainly developed with 64 bit MATLAB versions 2012b, 2014a, 2015a in Windows 7. When using SIMToolbox functions without the GUI, the MATLAB “Image Processing Toolbox” is required. SIMToolbox also requires the “MATLAB YAML” package to convert MATLAB objects to/from YAML file format. Note that this package is installed automatically when using the GUI.

## Availability of data

All raw and analyzed data is available on GigaDB at http://gigadb.org/site/index.

## Abbreviations

Av Int Proj: average intensity projection;
FOV: field of view;
H&E: hematoxylin and eosin;
ICE: Image Composite Editor;
MAP-SIM: maximum *a posteriori* probability SIM;
NA: numerical aperture;
LCOS: liquid crystal on silicon;
PSDca: circularly averaged power spectral density;
SIM: structured illumination microscopy;
WF: wide field.

## Ethics approval and consent to participate

Not applicable

## Consent for publication

Not applicable

## Author Contributions

KJ: acquired data, analyzed data, wrote the paper

GH: conceived project, acquired data, analyzed data, supervised research, wrote the paper

## Supplementary Information

### 1. Edge handling method during blurring step

Creating an estimate of the vignette profile is a crucial step of our devignetting process. While the average intensity projection removes most foreground information from the images, some coarseness remains in the vignette estimation after this step (Fig. S1a), especially for stitches with fewer than 50 tiles. The blurring step serves to eliminate only this non-vignette information and must preserve the illumination profile. As shown in Fig. S1b, use of the default Gaussian blurring function in ImageJ introduces a bright glow near to borders of the image, a significant artifact which does not reflect the original vignetting profile. Use of the ‘border limited mean’ filter, on the other hand, does not introduce this aberration, as shown in Fig. S1c.

**Figure S1:**
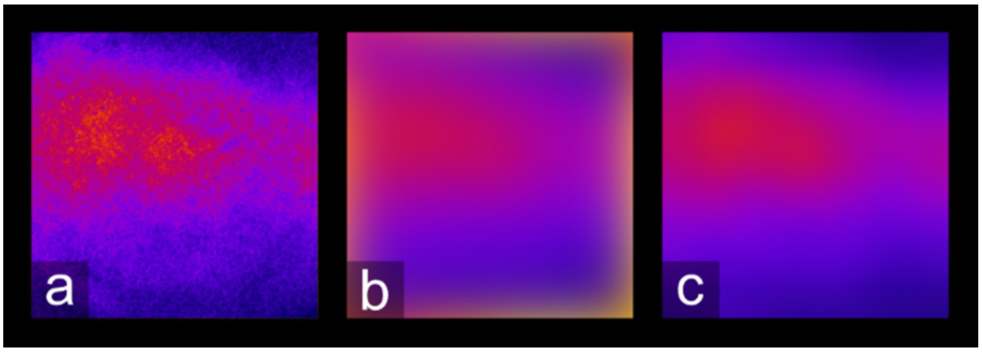
Edge handling during the blurring process. (a) is the result after performing an average intensity projection on a set of MAP-SIM tiles. (b) shows (a) after being blurred using ImageJ’s ‘Gaussian blur’ with a radius of 200 pixels, while (c) shows (a) blurred using the ‘border limited mean’ filter.

### 2. Example of data re-use: Analysis of periodic stitching artifacts in the frequency domain

Gone uncorrected, vignetting in the image tiles used to stitch a larger image can cause a noticeable grid pattern in the final stitched image. While readily noticeable upon viewing of the image, quantification of this pattern is useful for evaluation of methods which remove it. Since this stitching artifact arises from an illumination profile common to each tile, the period of this pattern in the stitched image can simply be represented by the spacing between tiles used during acquisition:

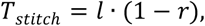

where *l* is the image width and *r* is the proportional overlap between image tiles. The parameters for the dataset visualized in Fig. S2 are *l* = 2048 px and *r* = 0.2, thus *T*_*stitch*_ = 1638 px, or ∼355 µm (as the dataset has a pixel size of ∼216.7 nm). In our setup, the camera sensor is square, so *T*_*stitch*_ is the same both the horizontally and vertically in the final image. As a pattern with a very consistent period, this grid artifact manifests in the FFT of an uncorrected stitch as series of bright peaks. As shown in Fig S2e, the location of the peaks corresponding to the fundamental frequency of the grid pattern agrees very well with the calculated *T*_*stitch*_. Figs. S2d and S2e show that the FFT of a properly corrected image contains no trace of the peaks evident in the uncorrected image.

**Figure S2:**
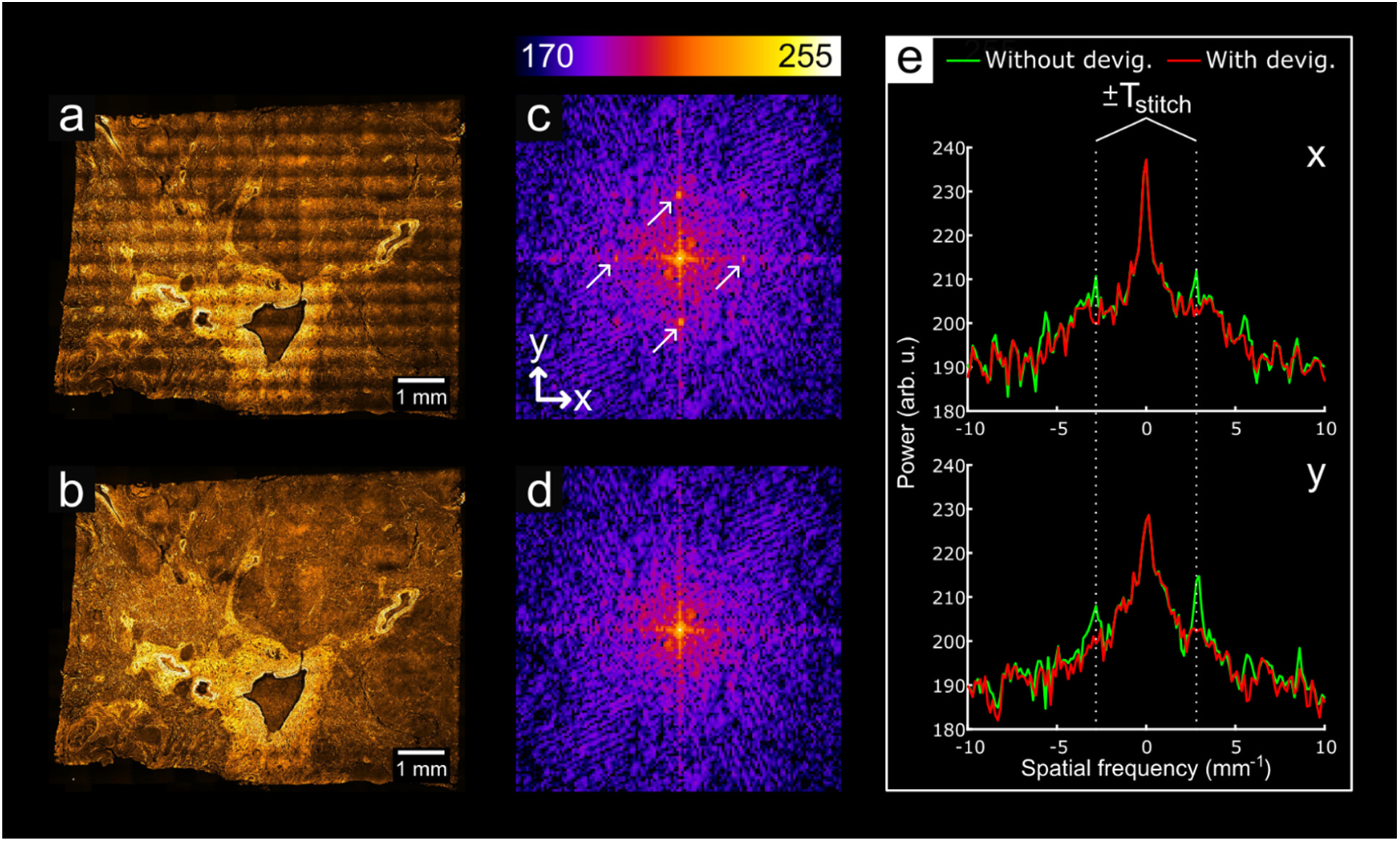
Analysis of periodic stitching artifacts in the frequency domain. (a) shows a stitched image where devignetting has not been applied to the tiles, while (b) shows a stitch of devignetted data. (c) and (d) show the central region of the FFT’s of (a) and (c), respectively. The arrows in (c) point to the fundamental peaks of the grid artifact. (e) shows plots of (c) and (d) through the center of each image along the x and y axes. The calculated value for *T*_*stitch*_ aligns very well with the peaks along both axes.

### 3. Multicolor, 100× stitched image using MAP-SIM

To demonstrate that the methods described in the main paper are not limited to medium-magnification, monochrome imaging of H&E stained histopathological slides, we used the same process to create a dual-color stitched image of fixed bovine pulmonary artery endothelial (BPAE) cells. While MAP-SIM does not require multiple angles of the illumination pattern, it can nonetheless process data acquired with such patterns. To demonstrate this, we used an illumination pattern with four different pattern angles for this acquisition. SIM reconstructions were performed on the acquired data, and the resulting tiles were stitched as described in the main paper. The imaging parameters for this dataset are detailed in Table S1, and the resulting images are shown in Figure S3.

**Table S1.** Imaging parameters for the data shown in Figure S3.

| Sample | Source company and product no. | SIM pattern no. of angles/phases | Exposure time, ms (illumination wavelength, nm) | No. of tiles | Objective mag./NA | Acquisition time, s |
| --- | --- | --- | --- | --- | --- | --- |
| BPAE cells | Thermo Fisher, F36924 | 4/34 | 200 (575 nm)<br>500 (470 nm) | 4 × 4 | 100×/1.35 | 505 |

**Figure S3.**
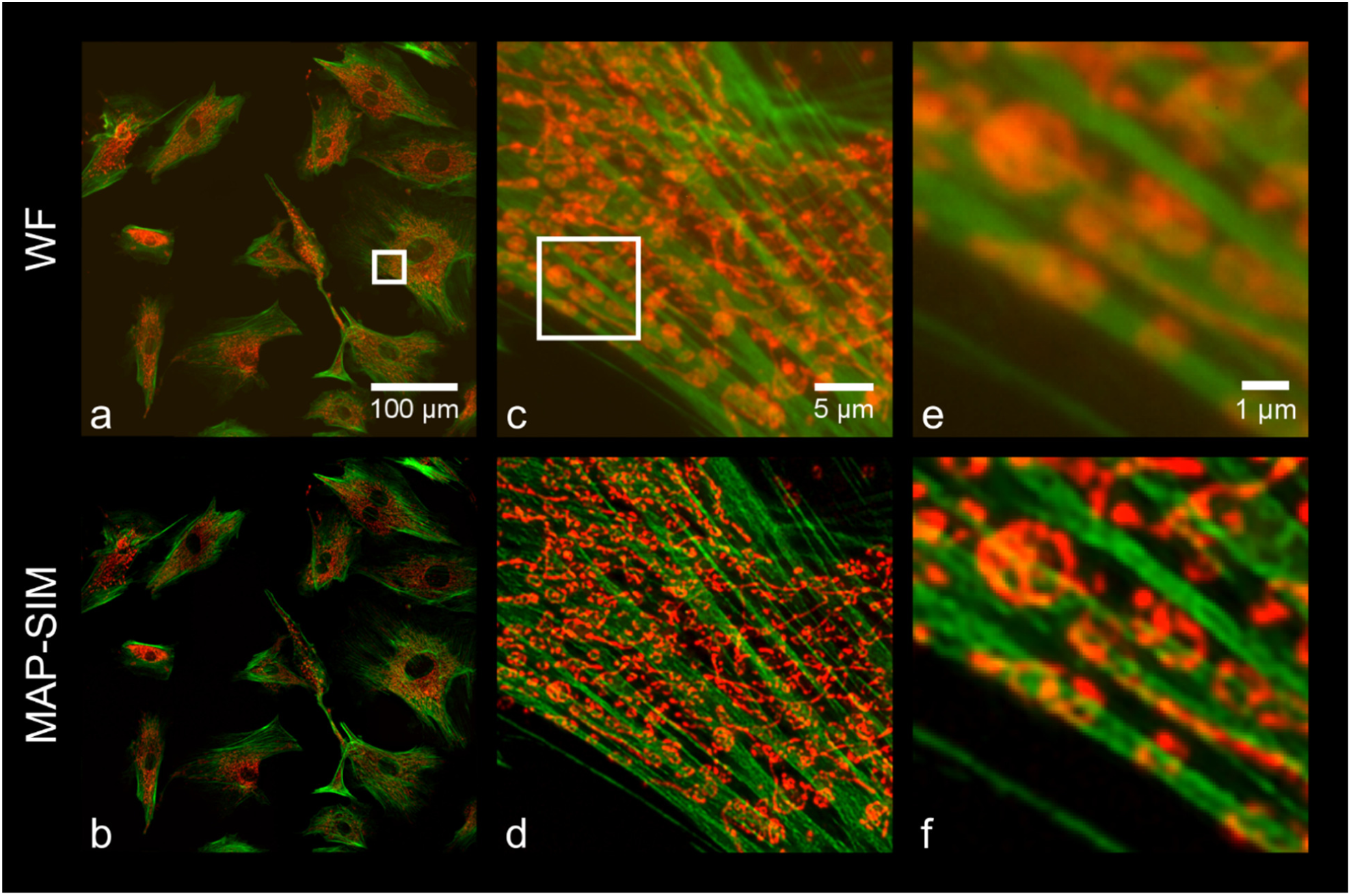
High magnification, dual-color, stitched SIM images of fixed BPAE cells. (a) and (b) show the full stitched image of BPAE cells, where Alexa Fluor 488 (actin) is shown in green and MitoTracker CMXRos (mitochondria) is shown in red. (c) and (d) show the region indicated in (a) for each reconstruction method, while (e) and (f) show the region indicated in (c).

### 4. Motorized stage and illumination triggering connection diagram

Another important aspect of the setup described in the main paper is the electronic synchronization between camera acquisition, stage sample movement, and illumination. As shown in Figure S4, the Andor IQ software serves as the hub for all signals in our system.

**Figure S4.**
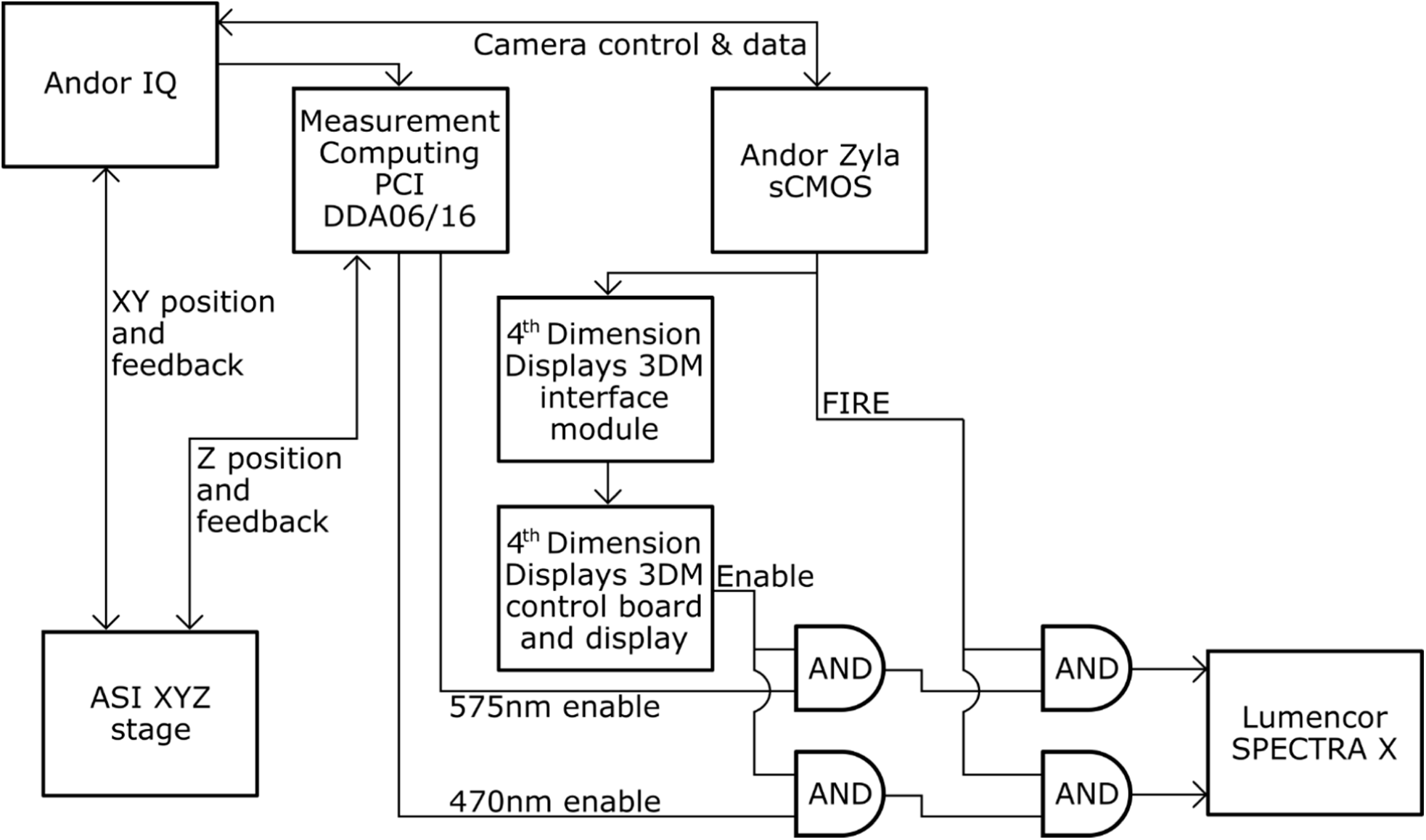
Simplified connection diagram.

While the Zyla camera and ASI XYZ stage receive signals directly from Andor IQ and the Measurement Computing PCI card, the illumination signals which IQ generates must be altered significantly before being sent to the light source. Firstly, the 4^th^ Dimension Displays (4DD) spatial light modulator (SLM) used in our setup will not produce an image on the sample if illuminated with a constant light source. Rather, a meaningful illumination pattern will only form if the light source is synchronized with an enable signal output from the 4DD control board. Therefore, the channel signals output by IQ are first modulated with the 4DD enable signal (this is performed by the leftmost AND gates pictured in Figure S4). Additionally, to reduce unnecessary light exposure to the sample, the light source is shut off whenever the camera sensor is not being exposed. This is accomplished simply by performing another logical AND of the result of the previous AND with the ‘FIRE’ signal output from the Zyla camera, as pictured in Figure S5.

**Figure S5.**
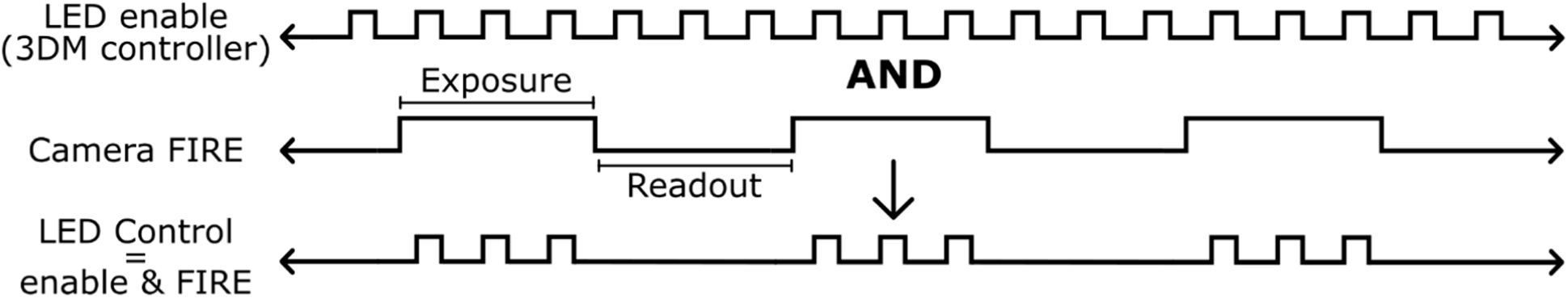
Illumination trigger signal logic. This logical operation occurs for each illumination wavelength at the rightmost AND gates pictured in Fig. S4.

